# Unveiling the Intercompartmental Signaling Axis: Mitochondrial to ER Stress Response (MERSR) and its Impact on Proteostasis

**DOI:** 10.1101/2023.09.07.556674

**Authors:** Jeson J Li, Nan Xin, Chunxia Yang, Larissa A Tavizon, Ruth Hong, Jina Park, Travis I Moore, Rebecca George Tharyan, Adam Antebi, Hyun-Eui Kim

## Abstract

Maintaining protein homeostasis is essential for cellular health. Our previous research uncovered a cross-compartmental Mitochondrial to Cytosolic Stress Response, activated by the perturbation of mitochondrial proteostasis, which ultimately results in the improvement of proteostasis in the cytosol. Here, we found that this signaling axis also influences the unfolded protein response of the endoplasmic reticulum (UPR^ER^), suggesting the presence of a Mitochondria to ER Stress Response (MERSR). During MERSR, the IRE1 branch of UPR^ER^ is inhibited, introducing a previously unknown regulatory component of MCSR. Moreover, proteostasis is enhanced through the upregulation of the PERK-eIF2α signaling pathway, increasing phosphorylation of eIF2α and improving the ER’s ability to handle proteostasis. MERSR activation in both polyglutamine and amyloid-beta peptide-expressing *C. elegans* disease models also led to improvement in both aggregate burden and overall disease outcome. These findings shed light on the coordination between the mitochondria and the ER in maintaining cellular proteostasis and provide further evidence for the importance of intercompartmental signaling.

## Introduction

A cornerstone of maintaining physiological health is the ability of our cells to maintain protein homeostasis (proteostasis) during challenging times. Our cells have developed unique defense mechanisms to detect and resolve proteotoxic stress through the activation of complex signaling pathways. These pathways mediate the upregulation of chaperone proteins to aid in the proper folding of the misfolded proteins in addition to reducing protein load by suppressing global protein translation ^1^. Throughout evolution, our cells have evolved to compartmentalize, yielding functionally unique spaces to enhance the efficiency of biological functions ^2^. With this also came the rise of compartmental stress responses within the ER, mitochondria, and cytosol. For quite some time, these cellular stress responses have been thought to be stand-alone systems, each encompassing its own unique set of regulatory elements and chaperone proteins. However, recent findings suggest the presence of a regulatory network that mediates an intercompartmental signaling axis.

Previously, we discovered that perturbation of the mitochondrial UPR (UPR^MT^) through RNAi targeting mitochondrial HSP70 (mtHSP70, HSP-6) in *C. elegans* resulted in an increase in the cytosolic heat shock response. This mitochondrial to cytosolic stress response (MCSR) improved cytosolic proteostasis in both *C. elegans* and human cell lines. Further evidence suggested that MCSR is regulated through the alteration of lipid metabolism, specifically ceramide and cardiolipin ^3^. Although this was the first evidence of interaction between the mitochondria and the cytosol mediated through *mtHSP70*, previous interactions between the mitochondria and ER have been described. Rizzuto *et al*. first characterized the importance of mitochondria and ER contact sites (MERCS) in the regulation of calcium signaling ^4^. MERCS has since been well studied and its role in multiple biological functions including Ca^2+^ signaling ^5^, lipid transport ^6^, mitochondria dynamics ^7^, and stress response ^8^ has been characterized. MERCS has also been implicated in an array of diseases including neurodegenerative disease ^9^ and cancer ^9,10^ and is currently being explored as a possible therapeutic target ^11^.

The ER is the major site of secretory protein synthesis, and the proper function of the UPR^ER^ is crucial to maintaining proper cellular function ^12^. While *mtHSP70* knockdown is able to induce MCSR and offers insight into the intercompartmental signaling axis between the mitochondria and cytosol, the question remains whether this axis exerts a dualistic influence over the ER. Indeed, our data suggests that the knockdown of *hsp-6* modulates UPR^ER^ by potentiating UPR^ER^-specific signaling pathways. Using animals harboring a transcriptional reporter of UPR^ER^ (*hsp-4*p::GFP), we observed a decrease in GFP fluorescence when treating the worms with tunicamycin followed by *hsp-6* knockdown. Recent evidence suggests a regulatory roles of mitochondrial proteins on the UPR^ER^ and Integrated Stress Response (ISR) ^13^ ^14,15,16^; however, the influence of the mitochondria on the UPR^ER^ is currently a field that has not been well studied. Hence, more extensive work is needed to fully characterize this signaling axis.

We have previously shown that *hsp-6* knockdown reduces polyglutamine aggregates in a *C. elegans* strain that expresses polyglutamine repeats within body wall muscles^3^. We hypothesize that this decrease in aggregate abundance occurs partially through the activation of the UPR^ER^ signaling pathways. The activation of the UPR^ER^ signaling pathway increases the protein folding capacity of the ER, which compensates for the increase in proteomic stress and ultimately helps to achieve ER stress resolution. When we perturbed the UPR^ER^ signaling pathway during *hsp-6* knockdown, we surprisingly found that the IRE1 branch of UPR^ER^ was inhibited. However, we found a significant increase in PERK-dependent eIF2alpha (eIF2α) phosphorylation, suggesting a decrease in global protein translation levels. Concomitantly, ER protein secretion capacity was also shown to be downregulated, suggesting a functional inhibition of the ER to further prevent the increase in misfolded proteins. Taken together, we infer that MERSR is able to increase proteostasis through dampening of IRE1 branch of UPR^ER^ signaling pathways and is able to further reduce proteomic stress by suppressing ER protein secretion, ultimately leading to the increase in the capacity of the ER to combat increasing proteostasis.

## Results

### Tunicamycin-induced ER stress is suppressed through the activation of MERSR

To determine the effect of *mtHSP70* knockdown on UPR^ER^, we utilized a *C. elegans* UPR^ER^ reporter strain where a GFP tag has been fused to the promoter of a UPR^ER^ chaperone (SJ4005, *hsp-4p*::GFP). Tunicamycin treatment of the worms induced ER stress, resulting in GFP fluorescence that can be measured and quantified using fluorescent microscopy. ER stress reporter worms were treated with tunicamycin during day 1 of adulthood followed by transfer onto RNAi plates containing *mtHSP70* targeting RNAi sequences. Loss of *mtHSP70* resulted in a reduction of GFP fluorescence post-tunicamycin treatment when compared to empty vector control (Figure 1a). To determine if the UPR^ER^ suppression by *mtHSP70* knockdown was limited to tunicamycin, we targeted 2 additional distinct UPR^ER^ activators. 1) NFYB-1 is a histone-like transcription factor that regulates mitochondrial function. Interestingly, it was found that NFYB-1 knockout exhibited cross-compartmental regulation and activation of UPR^ER^ ^17^. 2) VCP is an AAATPase that among many other things, regulates ER-associated degradation (ERAD) ^18^. The knockout of VCP is known to induce UPR^ER^ due to increased proteomic stress as a result of the increase in misfolded proteins ^19^. Ultimately, *mtHSP70* knockdown was able to reduce the UPR^ER^ induced by loss of either NFYB-1 (Figure 1b) or CDC-48, a *C. elegans* homolog of VCP (Figure 1c), suggesting that downregulation of *mtHSP70* reflects an intrinsic mechanism that regulates an intercompartmental UPR signaling axis during times of stress. Interestingly, when we target transcription factors involved in the UPR^MT^ signaling pathway, such as *dve-1*, a similar reduction in UPR^ER^ reporter fluorescence level is observed (Figure S1a), suggesting that the reduced UPR^ER^ activation is a direct consequence of alterations that originate within the mitochondria. Mitochondrial stress activated through *mrps-5*, *spg-7* or *cco-1* knockdown was unable to reduce UPR^ER^ signaling during tunicamycin treatment (Figure S1b) or in *nfyb-1* mutants (Figure S1c), suggesting that *mtHSP70* knockdown triggers a distinct mitochondrial stress that elicits a signaling cascade between the mitochondria and the ER. Collectively, our data suggests that *mtHSP70* knockdown not only triggers MCSR, but also triggers a cross-compartmental signaling axis, activating a mitochondrial to ER Stress Response.

**Figure 1.**
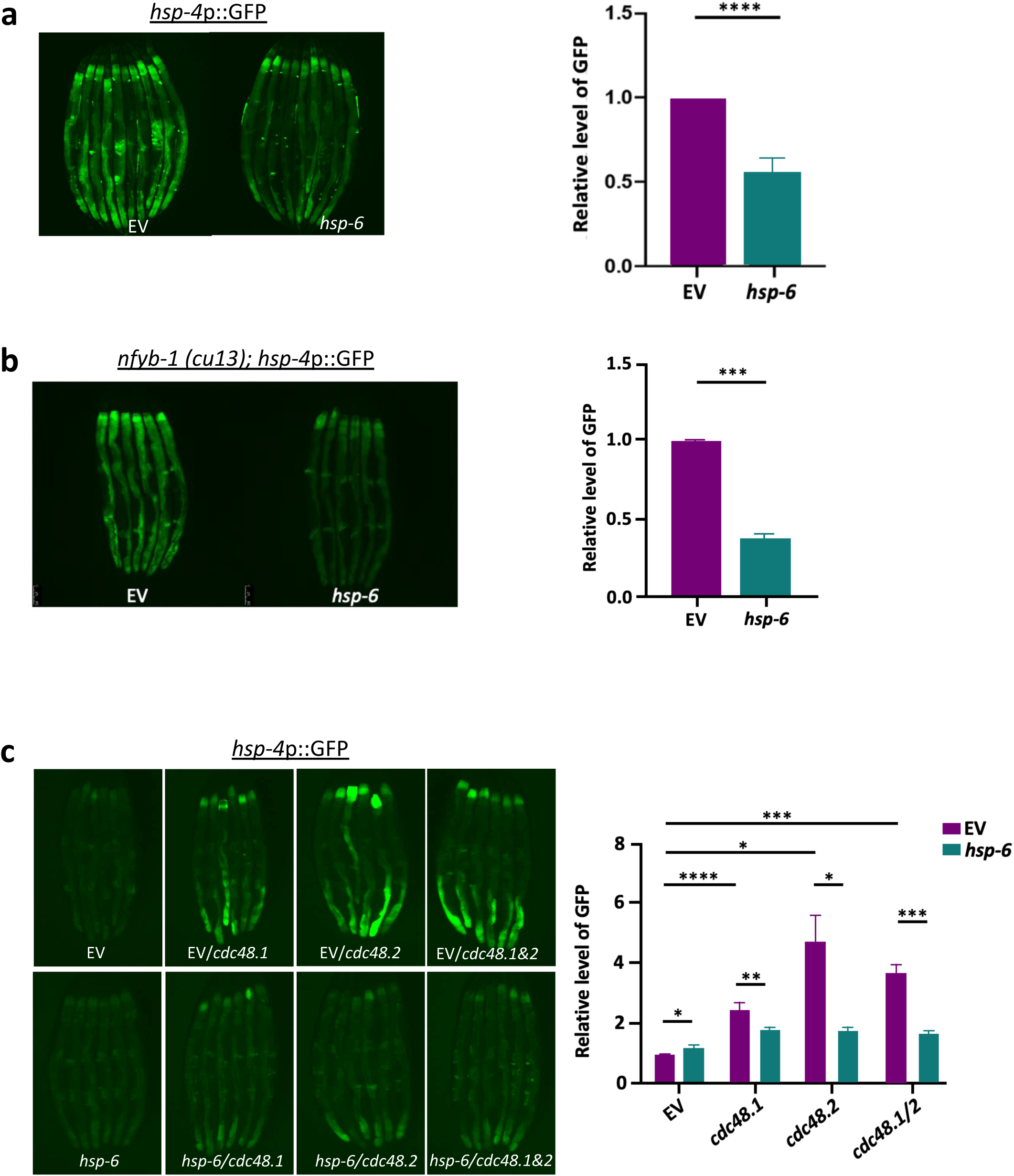
Inhibition of *hsp-6* results in the reduction of ER stress response. *hsp-4p*::GFP, the UPR^ER^ reporter animals were transferred to the RNAi-containing plates L4 adulthood (55 hours post bleaching). On day 1 adulthood (72 hours post bleaching), animals were treated with **a)** tunicamycin for 4 hours in order to induce ER stress. Following tunicamycin treatment, the worms were transferred onto RNAi-containing plates and grown at 20°C until day 3 of adulthood and were imaged to assess ER stress levels. Other forms of ER stress, including **b)**nfyb-1 (cu13) mutant and **c)** VCP (*cdc-48.1* and *cdc-48.2*) knockdown, were tested in the same manner to further validate that our observation was not tunicamycin specific but rather a conserved physiological mechanism. When multiple RNAi treatments were needed, the same amount of dsRNA-expressing bacteria were mixed. The RNAi target genes are as indicated, EV: empty vector control. The graphs show the mean+/-SD of the animals with representative images of animals, n>=6 (three biological repeats). Each RNAi-treated cohort was compared to the EV control to compare GFP induction.

### Lipid metabolism mediates the ER suppression by *hsp-6* RNAi

We previously observed that MCSR is mediated through altered lipid metabolism, in particular, the downregulation of ceramide and the upregulation of cardiolipins ^3^. In *C. elegans*, *de novo* ceramide synthesis depends on the function of three separate ceramide synthase enzymes HYL-1, HYL-2 and LAGR-1, and the knockdown of these enzymes can alter the global sphingolipid landscape of the animals ^20^. To investigate the possibility that sphingolipid metabolism also regulates MERSR, we targeted several critical enzymes in the *de novo* sphingolipid synthesis pathway in addition to cardiolipin synthase (*crls-1*) through RNAi. By inducing UPR^ER^ in ER stress reporter worms with tunicamycin treatment followed by the perturbation of key sphingolipid synthesis enzymes through RNAi, we found that the knockdown of ceramide synthases exhibited a decrease in UPR^ER^ reporter intensity similar to that of *mtHSP70* RNAi (Figure 2a). Other enzymatic targets within the *de novo* synthesis pathway, however, did not yield such results (Figure 2b). Conversely, cardiolipin synthase (*crls-1*) knockdown negated the inhibition of UPR^ER^ through *mtHSP70* knockdown (Figure 2c). These results suggest that not only do ceramide and cardiolipin mediate the stress signaling response between the mitochondria and the cytosol, but they also regulate the UPR^ER^ response initiated through the knockdown of *mtHSP70.* The decrease in UPR^ER^ reporter signaling due to the knockdown of ceramide synthase could be the result of increased UPR^ER^ threshold caused by changes in ceramide levels, enhancing the capability of the cells to combat protein stress and, therefore, reducing UPR^ER^ ^21^. Furthermore, *nfyb-1* deleted mutant animals constitutively activate higher extent of UPR^ER^ (Figure S1c), which could be reduced through the activation of MERSR (Figure 1b). Interestingly, the lipid profiles of these *nfyb-1* mutants exhibit a high level of ceramide and low levels of cardiolipin compared to their wild-type counterparts ^17^. The reduction in UPR^ER^ observed in these mutant animals through MERSR activation could be the result of reversing the ceramide and cardiolipin content, reducing ceramide while increasing cardiolipins to reduce lipid signals activating UPR^ER^.

**Figure 2.**
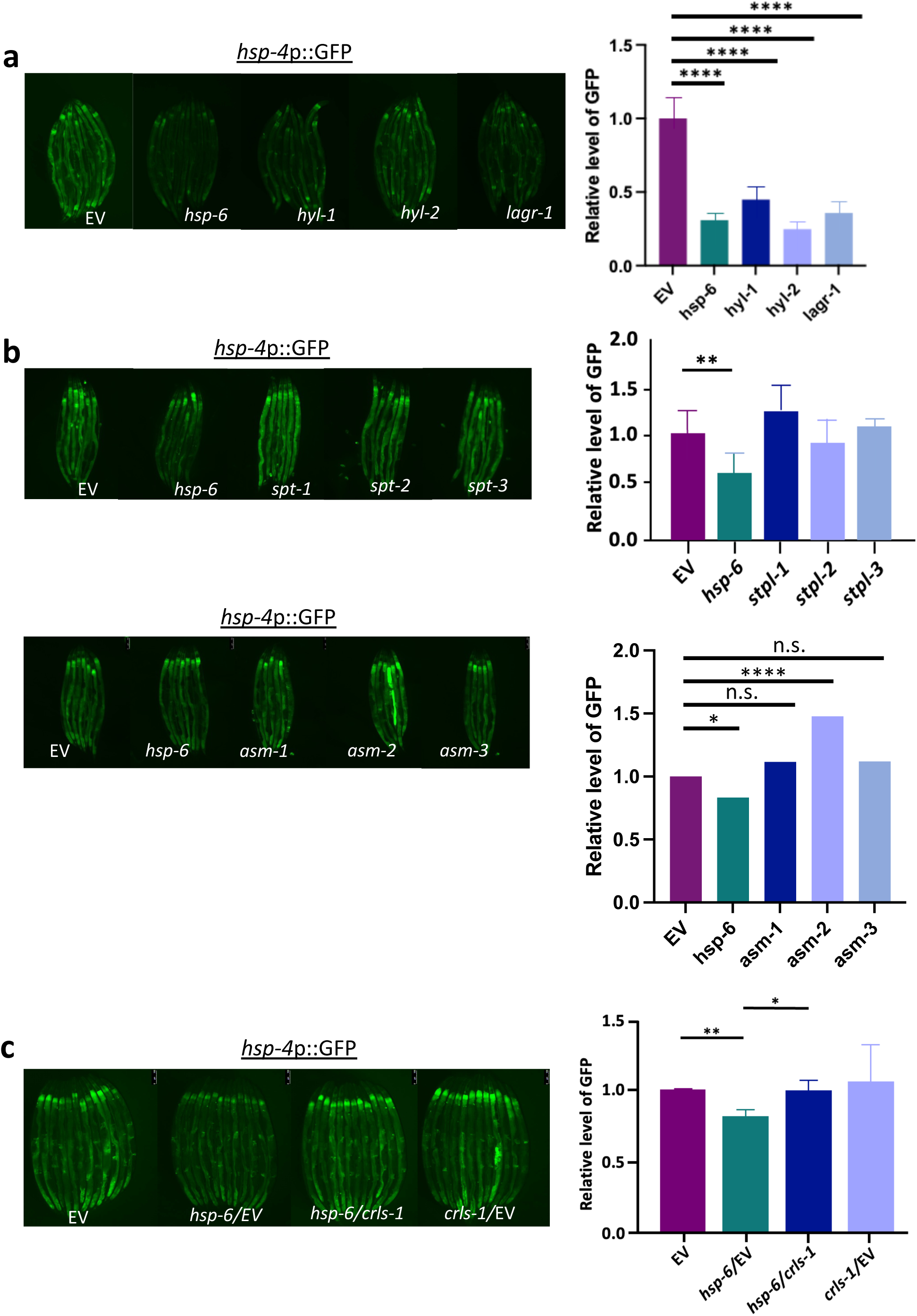
The reduction of ER stress by *hsp-6* RNAi is partially regulated through lipid metabolism. a) *hsp-4p*::GFP, the UPR^ER^ reporter animals were used to determine the effects of *hyl-1*, *hyl-2* and *lagr-1* on UPR^ER^ induction as described in Figure 1. b) Other sphingolipid enzymes along the *de novo* synthesis pathway and the salvage pathways were also tested in the same manner. c) Knockdown of cardiolipin synthetase *crls-1* positively regu^la^ted induction of UPR^ER^. When multiple RNAi treatments were needed, the same amount of dsRNA-expressing bacteria were mixed. The RNAi target genes are as indicated, EV: empty vector control. The graphs show the mean+/-SD of the animals with representative images of animals, n>=6 (three biological repeats). Each RNAi-treated cohort was compared to the EV control to compare GFP induction.

### MERSR is mediated through the UPR^ER^ signaling pathway

UPR^ER^ signaling is mediated through three separate stress sensory pathways that work together to increase the capacity of the ER to combat proteomic stress ^1^. During proteotoxic stress, membrane IRE1 oligomerizes, initiating splicing of XBP-1 mRNA to the active transcription factor XBP-1s, resulting in the upregulation of various genes, including major chaperone proteins in order to aid in the proper folding and recycling of misfolded proteins ^22^ ^23^. In addition to increasing chaperone levels, another major compensatory mechanism of the ER is the PERK-eIF2α signaling pathway, which attenuates global protein translation levels and reduces the total protein load of the ER ^24^. To determine the effects of MERSR on UPR^ER^ signaling pathways, we perturbed the UPR^ER^ signaling pathway during *mtHSP70* knockdown and found that the double knockdown of *mtHSP70* with *ire-1* or *xbp-1* resulted in the loss of MERSR activation (Figure 3a). Proper UPR^ER^ signaling is reliant on the dimer/oligomerization of IRE1, which initiates splicing of *xbp-1* mRNA. To investigate the impact of *mtHSP70* knockdown on ectopic UPR^ER^ activation at two different points within the same pathway, we used 1) overexpression of IRE1 lacking the luminal domain, which activates UPR^ER^ in the absence of proteotoxic stress but still requires proper dimer/oligomerization and 2) overexpression of *xbp-1*s (active form of *xbp-1*), allowing to bypass IRE1 dimer/oligomerization and activate UPR^ER^ (Figure 3b). Overexpression of IRE1 lacking the luminal domain (*ire-1a* (344-967aa)) in the intestine induced UPR^ER^ reporter ^25^. This induction was moderately reduced upon *mtHSP70* knockdown while *mtHSP70* knockdown had no impact on *xbp-1*s overexpression-induced UPR^ER^ (Figure 3c and d). This suggests that the suppression of UPR^ER^ occurs upstream of *xbp-1*, possibly at a membrane level where IRE1 resides. One possibility is that *mtHSP70* knockdown leads to an alteration in membrane fluidity, compromising the dimerization of the UPR^ER^ sensor IRE1. This is further corroborated by recent findings that the knockdown of ER-resident proteins alters the ER membrane lipid content, increasing mitochondria and ER contact through increasing membrane order ^26^. Furthermore, alterations in sphingolipid, including ceramide content within the ER membrane, have been shown to induce UPR^ER^ in the absence of aberrant proteostasis ^27^, providing further evidence that membrane lipid content can directly influence the UPR^ER^ signaling pathways.

**Figure 3.**
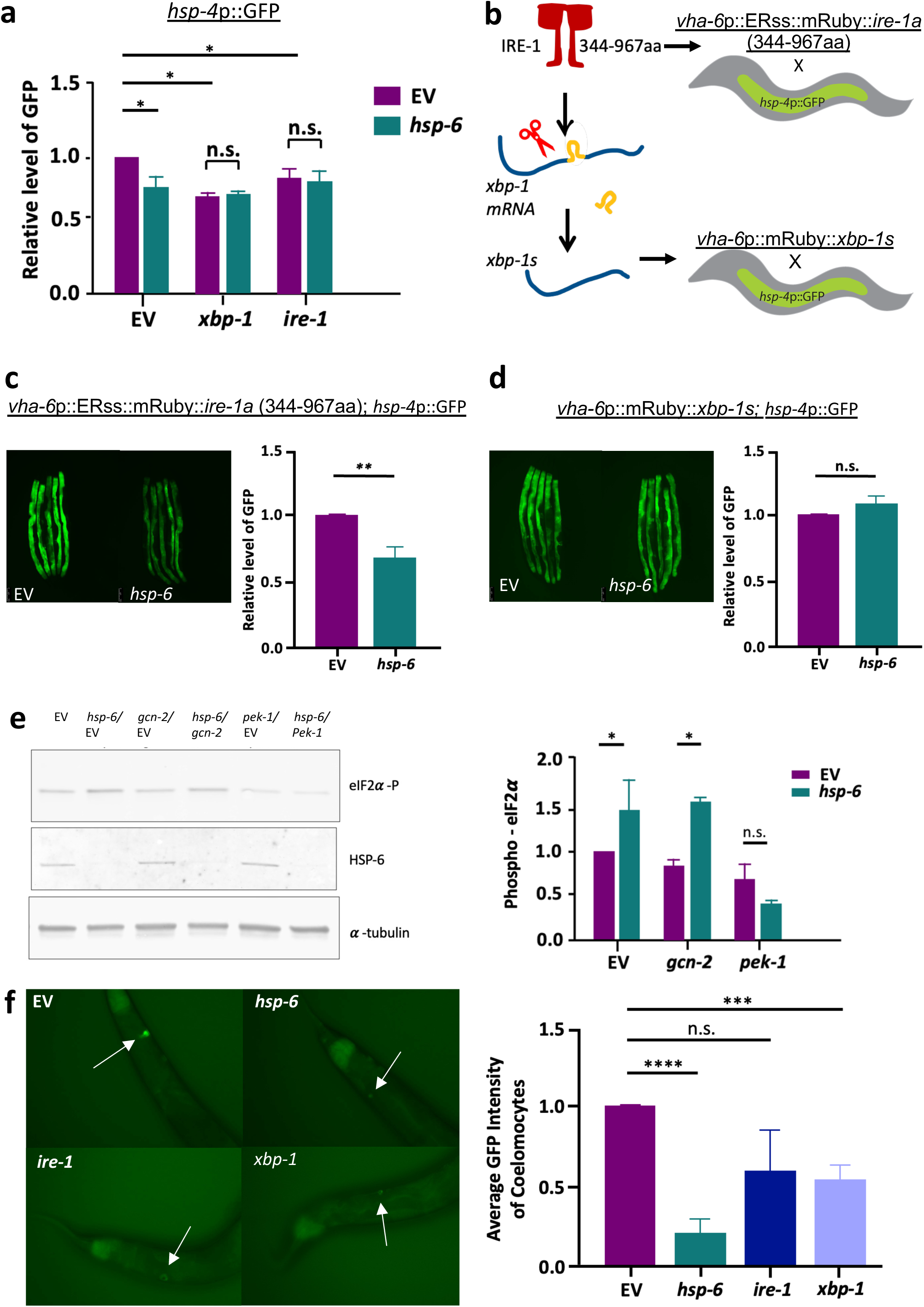
The reduction of ER stress by *hsp-6* RNAi is the result of IRE-1 and PERK modulation. **a)** Tunicamycin-treated animals induced UPRER in an *xbp-1* and *ire-1* dependent manner. **b)** *hsp-4p*::GFP, the UPR^ER^ reporter strain was crossed with animals that constitutively activate UPR^ER^ through the IRE-1-XBP-1 pathway. Animals expressing constitutively active IRE-1a (344-967aa) that are lacking part of the luminal domain and animals expressing *xbp-1s* both activate UPR^ER^ (c and d). **c and d)** *hsp-6* RNAi reduced UPR^ER^ activation only in the animals expressing constitutively active IRE-1a (c) but not in the animals expressing the *xbp-1* spliced form, *xbp-1s*. MCSR regulates UPR^ER^ at the IRE-1 level on the ER membrane. Animals were transferred to *hsp-6* RNAi plates or the empty vector control plates on late L4/early adult and were imaged on day 3 (72 hours of RNAi treatment). ***e*)** Wild-type N2 animals were treated with EV, *gcn-2*, or *pek-1* RNAi from hatch to the late L4 stage, before transferring to double RNAi treatment or to continue single RNAi treatment. Two individual double RNAi treatment groups were set up utilizing a 1:1 ratio of EV (purple) or *hsp-6* (teal) combined with *gcn-2* or *pek-1* for overnight treatment. *Hsp-6* RNAi increased the eIF2α phosphorylation in a *pek-1-*dependent manner. **f)** DAF-28::GFP transgenic animals were treated with indicated RNAi, or the empty vector control from day 1 of adulthood. Animals were imaged for coelomocyte GFP content on day 4. The graph shows mean+/-SD of GFP intensity normalized to empty vector control, n>=6 with three biological repeats.

During times of proteotoxic stress, the activation of the UPR^ER^ have also been shown to decrease global protein load through the phosphorylation of PERK-eIF2α signaling cascade ^28^. Thus, we examined the levels of eIF2α phosphorylation during *mtHSP70* knockdown. Along with PERK, we also targeted *gcn-2*, a PERK-independent eIF2α kinase activated primarily through amino acid starvation ^29^ and reactive oxygen species produced during mitochondrial stress ^14^. Our data shows that *mtHSP70* knockdown enhances the phosphorylation level of eIF2α (Figure 3e), while the knockdown of *gcn-2* had no effect on either the induction or the suppression of UPR^ER^ upon tunicamycin treatment (Figure S2a). This suggests that MERSR is able to increase the phosphorylation levels of eIF2α in a PERK-dependent manner. Moreover, when we performed a double knockdown of *mtHSP70* and *pek-1*, the inhibition of UPR^ER^ was no longer observed (Figure S2a), suggesting that the activation of MERSR is PERK-dependent. PERK may also contribute to the repression of XBP1 targets ^30^. Altogether, our data suggest that *mtHSP70* knockdown enhances proteostasis in the ER through the increase of eIF2α phosphorylation via PERK signaling, reducing global protein translation levels to compensate for increasing ER stress.

Given that MERSR reduces global protein translation through the mediation of UPR^ER^ signaling, we further assessed the function of the UPR^ER^ by investigating its secretory capacity regulated by the IRE1 signaling pathway. In *C. elegans*, DAF-28 is one of 40 insulin-like peptides ^31^. DAF28::GFP fusion protein is processed within the ER and secreted to the body cavities of the worms ^32^, eventually absorbed by the coelomocytes, phagocytic and macrophage-like cells that engulf surrounding materials for degradation ^33^. GFP signal within the coelomocytes can be used as a readout to determine the ER secretion capacity of the cells ^34^. We observed that the knockdown of *mtHSP70* greatly reduced the GFP fluorescence within the coelomocytes (Figure 3f) to a level similar to that of ER dysfunction induced through the knockdown of *ire-1* and *xbp-1*. Western blot analysis of both of the transcriptional and translational *daf-28* expression reporter worms further confirmed that the reduction observed in the coelomocytes during *mtHSP70* RNAi was the result of decreased secretion rather than changes in the innate expression of *daf-28* (Figure S2b). This data suggests that in addition to decreasing global translation through the upregulation of PERK-eIF2α signaling, *mtHSP70* knockdown also inhibits ER secretory functions. This reduction in ER protein secretion could serve as a compensatory mechanism during times of stress to further slow down the proteomic flux.

### MERSR and ceramide reduction can benefit both ER and Cytosolic Stress-Related Neurodegenerative Diseases

Dysfunctional ER stress is a common phenotype in many neurodegenerative diseases, including Alzheimer’s, Huntington’s, and Parkinson’s Disease ^35^. Early notions of ER stress denoted its function as deleterious ^36^; however, recent studies have suggested that the response of UPR^ER^ in neurodegenerative diseases is highly dynamic ^37^. Mild UPR^ER^ is upregulated during early stages of AD in order to resolve amyloid-β (Aβ) aggregate deposition, however, chronic Aβ burden as a result of advanced AD and constitutive UPR^ER^ activation triggers downstream apoptotic pathways, leading to cell death ^38^. Aβ metabolism is a constant cycle, starting from the synthesis of Amyloid Precursor Protein (APP) in the ER, followed by the export to the Golgi and ultimately to the plasma membrane where it is cleaved by beta-secretase and gamma-secretase to form mature Aβ ^39^. While an ortholog of APP containing beta-secretase sites is absent in *C. elegans*, several disease models expressing the disease-causing human Aβ (1-42) peptide have been generated. One of the earliest models of Aβ worms was thought to have been able to generate Aβ (1-42) peptides ^40^, but further research established that the final Aβ product was a truncated version (3-42) due to mis-cleavage of a synthetic signal peptide ^41^. However, by modifying the artificial signal peptide on the Aβ plasmid, worm strains that produce Aβ (1-42) peptide have now been generated ^42^. By taking advantage of one of these worm lines, we explored the impact of MERSR on Aβ aggregate burden in *C. elegans*. We hypothesized that the pro-proteostasis phenotype observed with MERSR could decrease the Aβ burden, leading to improvements in the overall disease state. To evaluate the physiological effects of the Aβ burden, neuronal RNAi-sensitive worms expressing pan-neuronal Aβ (1-42) were put under the effects of mild heat. At 25°C, the increased temperature accelerates the aggregate formation process, leading to rapid paralysis of the worms. However, the neuronal knockdown of *mtHSP70* is able to delay the rate of paralysis, suggesting a decrease in Aβ aggregate burden (Figure 4a). Similarly, the knockdown of ceramide synthases (*hyl-1* and *hyl-2)* also improved the paralysis of neuronal Aβ-expressing animals. In addition, both *hsp-6 and hyl-1* knockdown also improved the lifespan of these worms (Figure 4c & d), providing further evidence that lipid metabolism plays an integral role in the regulation of cross-compartmental stress response.

**Figure 4.**
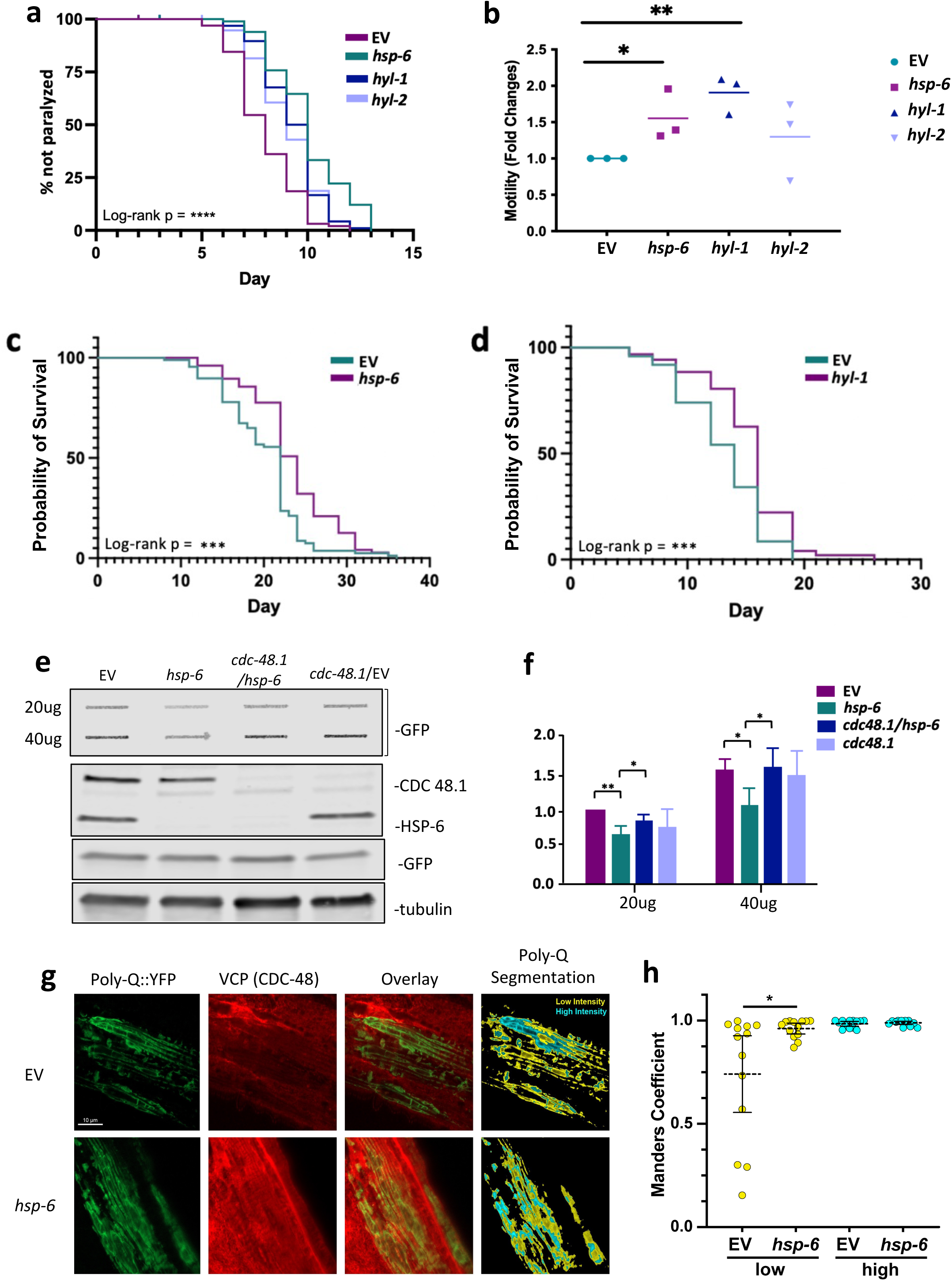
The knockdown of *hsp-6* reduces protein aggregation in poly-glutamine disease model worms. **a)** Paralysis of pan-neuronal Aβ expressing animals crossed with neuronal RNAi sensitive line (GRU102;TU3401). 100 animals per RNAi condition were incubated at 25°C to accelerate the production of the neuronal toxic aggregates, which led to paralysis. Treatment of RNAi against *hsp-6, hyl-1, hyl-2 and* EV were used to determine improvements on paralysis during heat shock. Log-rank P value was obtained comparing RNAi to empty vector control: P<0.0001 for all *hsp-6*, *hyl-1*, and *hyl-2*. **b)** Animals expressing polyglutamine aggregates (Q40::YFP) within the neurons (AM101) were crossed with a neuronal RNAi-sensitive worm line (TU3401). The resulting AM101;TU3401 showed decreased motility as the result of increased polyglutamine aggregate formation within the neurons. Motility was determined using a body bending assay that measures the number of body bends per 30 seconds. The relative motility was plotted by normalizing with empty vector control, and the rate was compared to that of the empty vector control (Three biological repeats with n>12, mean +/- SD). **c)** Lifespan of wildtype N2 worms treated with *hsp-6* RNAi. Animals were transferred to RNAi-containing plates at day 1 adult. P<0.001. **d)** Lifespan of the animals expressing Aβ in neurons with neuronal-specific RNAi treatment as shown in (a GRU102; TU3401). P<0.001. **e)** The accumulation of poly-glutamine aggregates is reduced upon MERSR. Transgenic worms expressing polyglutamine aggregates (Q35::YFP) in the body wall muscle (AM140) were treated with indicated RNAi or the empty vector control from day 1 adulthood. Animals were collected on day 5. Protein aggregates were measured by applying protein lysate onto a cellulose acetate membrane through a vacuum slot blotter. The membrane was blotted with GFP antibody to detect the Q35::YFP aggregates. RNAi knockdown of *hsp-6* results in less accumulation of Q35::YFP trapped on the membrane. Additional knockdown of *cdc-48.1* abolished the effect of *hsp-6* RNAi. 20ug of total protein lysate from each RNAi treatment was applied to SDS-PAGE followed by Western blot of alpha-tubulin, GFP, VCP, and HSP-6 antibody (lower panels). Total protein and total Q35::YFP protein levels were both at equivalent levels across all specimens. The VCP blot and HSP-6 blot showed that the RNAi knockdown of the targeted protein worked efficiently. **f)** Graph shows mean+-SD of filter trapped Q35::YFP from three biological repeats. **g)** Representative confocal image of polyQ::YFP (green) and VCP (CDC48, red) localization in the body wall muscle of AM140. Segmentation of polyQ::YFP by emission intensity (low, yellow; high, cyan). **h)** Colocalization of VCP with polyQ as measured by the Meanders correlation in high (yellow in the segmentation image) and low-intensity (cyan in the segmentation image) areas of polyglutamine (Poly-Q::YFP) in the empty vector control and *hsp-6* knockdown worms. Bars show mean ± 95% CI. Two-tailed Mann–Whitney tests show p=0.05 (*). n =13 (EV); n=14 (*hsp-6*).

Huntington’s Disease is characterized by the expansion of the polyglutamine tracks in mutant Huntingtin (htt) proteins, resulting in neurotoxic htt aggregate formation ^43^. The presence of the neurotoxic htt aggregates inhibits the ER-associated degradation pathway, resulting in increased proteomic stress and the activation of UPR^ER^ ^35^. To determine whether MERSR also improves the proteostasis of animals expressing neuronal polyglutamine repeats, we examined the animals expressing neuronal polyglutamine (Q40::YFP) repeats. Activation of MERSR through *mtHSP70* knockdown, or the knockdown of *hyl-1,* improved motility, evident in the increase in the number of body bends of the worms (Figure 4b). Similarly, the expression of polyglutamine (Q35::YFP) repeats in the body wall muscle showed decreased polyglutamine aggregation (Figures 4e and f). Interestingly, this decrease in polyglutamine aggregates required the presence of VCP (CDC-48 in *C. elegans*), an essential component in ER-associated Degradation (ERAD) and the Ubiquitous Proteasome System (UPS). This data suggests that a properly functioning ERAD system is required for the proper cellular processing of these neurotoxic aggregates. Further evidence suggests that VCP is highly correlated with densely packed polyglutamine aggregates, and upon MERSR activation, a portion of VCP becomes highly associated with loosely packed polyglutamine aggregates. After immunostaining VCP and polyglutamine aggregates in the body wall muscle, the aggregates were categorized into loosely packed (yellow) and densely packed (cyan) by the staining intensity (Figure 4g). In control animals, VCP is predominantly colocalized with densely packed aggregates. However, in *mtHSP70* RNAi-treated animals, there was a notable increase in VCP colocalization with loosely packed aggregates (Figures 4g and h). Consistent with this data, research has shown that mutant htt oligomers and not aggregates are responsible for the activation of UPR^ER^ ^44^. These data suggest that the induction of MERSR is able to mobilize VCP to a subset of polyglutamine aggregates. Although the downstream target of this VCP mobilization is unknown, given that the VCP function in aggregate handling is well characterized, we can postulate that VCP recruitment to the aggregates serves as a trigger for aggregate degradation through the UPS or macroautophagy. Finally, what would happen if MERSR can not reduce IRE1 activation in animals undergoing proteotoxicity? Animals overexpressing *xbp-1s* were crossed with animals expressing polyglutamine in body wall muscle and subjected to the lifespan assays (Figure S2c). These worms have short lifespan, and none of the RNAi treatments that ameliorate proteotoxicity were able to extend their lifespan. Interestingly, *hyl-2* RNAi further shortens its already short lifespan. This data suggests that uncontrolled UPR^ER^ activation does not improve the physiology of the animals and requires intricate regulation, particularly by MERSR in this study.

These data suggest that MERSR activation and ceramide perturbation through ceramide synthase knockdown can improve the disease outcome of both cytosolic- and ER-associated neurotoxic aggregate worm lines. Taken together, our findings suggest that the increase in mitochondrial stress triggered through *mtHSP70* knockout coordinates both cytosolic and ER stress responses, leading to enhanced cellular proteostasis through the upregulation of UPR^ER^ signaling pathways (Figure S3). Activation of MERSR in proteotoxicity models demonstrates its efficacy and inducibility in post-mitotic tissues such as neurons, further emphasizing the importance of this signaling pathway and the need for future research.

## Discussion

Here, we have identified an unexpected role of MCSR in promoting the balance of proteostasis by regulating the UPR^ER^ signaling pathways in *C. elegans*. Our study suggests that the knockdown of *mtHSP70* not only coordinates mitochondria to cytosolic stress response (3) but also elicits an intercompartmental signaling axis between the mitochondria and the ER. Mitochondrial stress triggered by the knockdown of *mtHSP70* results in decreased UPR^ER^ inducibility by multiple ER stress inducers, including tunicamycin, VCP knockout, and the loss of NFYB-1. The suppression of the IRE1 branch of the UPR^ER^ is mediated through perturbations of sphingolipid metabolism, suggesting the possibility of bioactive lipids serving as mediators along this intercompartmental signaling axis. The proper function of the UPR^ER^ is dependent on several signaling cascades. The PERK-eIF2α signaling cascade results in an increase in the phosphorylation of eIF2α, leading to a reduction in global protein translation levels. We have shown here that reduced UPR^ER^ inducibility is a direct result of increased PERK-dependent eIF2α phosphorylation in combination with decreased ER protein secretion.

Mitochondrial HSP70 (mtHSP70, mortalin, HSPA9, Grp75, HSP-6) is widely known for its beneficiary functions. It was first described as a substrate for protein translocation into the mitochondrial matrix ^45^. However, as research on the subject has increased, the role of mortalin has begun to expand. Mortalin reduces the formation of reactive oxygen species and lipid peroxidation, protecting the mitochondria from oxidative stress ^46^. In addition, mortalin exhibits protective properties against neurotoxicity in AD ^47^ and PD ^48^ through targeting of Aβ and alpha-synuclein, respectively. In light of this, an insightful review was recently published describing the link between mortalin and neurodegenerative disease ^49^. Interestingly, in our study, the post-developmental knockdown of *mtHSP70* in *C. elegans* conferred a protective phenotype against proteomic stress, suggesting a previously undiscovered role of mortalin. In support of our data, Honrath *et al*. also demonstrated a protective phenotype against glutamate-induced oxidative cell death in Grp75 knockout HT22 cells ^50^. However, this protection did not extend to drug-induced ER stress. This contradiction could be explained by the utilization of cancer cells in testing for ER stress protection. Cancer cells exhibit high levels of innate ER stress, and treatment with tunicamycin or thapsigargin may exacerbate ER stress and induce apoptotic signaling. In recent cancer research, mortalin has become a target for drug discovery due to its ability to bind P53 and prevent its nuclear localization ^51^. However, this colocalization is not observed in non-cancer cells ^52^, and the protective phenotype induced through *mtHSP70* knockdown is still unknown. Despite this, given the significant changes in lipid metabolism, specifically cardiolipin and ceramide, it is likely that *mtHSP70* knockdown-induced mitochondrial stress could result in the remodeling of the lipid composition within the mitochondrial membrane. Cardiolipin is a key regulator in the maintenance of mitochondrial membrane integrity and cristae morphology ^53^. Proper cardiolipin metabolism is also responsible for mediating proper protein-protein and protein-membrane interactions ^54^ in addition to stabilizing the respiratory chain complex ^55^. Although the mechanism of regulating cardiolipin to ceramide ratio between the mitochondria and the ER during *mtHSP70* knockdown remains undetermined, recent evidence revealed that *mtHSP70* is inserted into the mitochondrial membrane and possesses an affinity for negatively charged phospholipids such as cardiolipin ^56^. Indeed, cardiolipins are externalized during mitochondrial stress and serve as a signal for the induction signaling for mitophagic ^57^ and apoptotic ^58^ signaling; however, the exact mechanism of how cardiolipins regulate cross-compartmental signaling pathways will require further research.

Furthermore, we have also shown that MERSR induction also improved disease states in our Aβ models. Although the Aβ aggregates we observe in our worm models are primarily cytosolic ^63^, the expression of Aβ (1-42) requires the cleavage of a synthetic signaling peptide, which occurs at the ER ^64^ ^42^. We have shown that MERSR reduces ER protein secretion. Therefore, it would be prudent to suggest that the cleavage of Aβ may be compromised during MERSR induction, which would result in the decrease in disease burden that we observed in our Aβ worm model. Interestingly, ERAD-related chaperone VCP was associated with loosely packed polyglutamine aggregates in the muscle of AM140 animals when MERSR was activated. In these animals, MERSR reduced protein aggregates in a VCP-dependent manner, and the knockdown of VCP resulted in the loss of proteostasis improvement. Although during ERAD, VCP is recruited to the surface of the ER by ubiquitin ligases to aid in the clearance of misfolded proteins ^59^, recent evidence shows strong support for the colocalization of VCP with neurotoxic aggregates ^60^. The regulatory role of VCP in protein homeostasis occurs at multiple levels. During the initial process of aggregate formation, VCP regulates the UPS to detect and ultimately degrade misfolded proteins ^61^. However, as aggregate burden increases, VCP is able to regulate macroautophagy and aid in the removal of excess protein aggregates and damaged organelles to restore homeostasis ^62^. In our study, we showed that VCP is essential in reducing polyglutamine aggregates during MERSR; however, the downstream mechanisms of how aggregate clearance is achieved were beyond the scope of this paper. This does leave us with an exciting future research question of how mitochondrial stress regulates VCP activity to improve proteomic stress. Recent evidence has already shown VCP’s involvement in the regulation of mitochondria and ER contact ^65^ ^66^, further exacerbating the need for elucidation in this subject.

In conclusion, we have demonstrated for the first time, within our knowledge, the capabilities of a mitochondrial chaperone protein to regulate UPR^ER^ signaling through the perturbation of lipid metabolism. Current understanding of the relationship between the mitochondria and the ER is incomplete, although research on this subject has increased due to advances in biochemical and fluorescent microscopy. We know that mitochondria and the ER form contact sites in order to modulate membrane lipid composition and proper Ca^2+^ homeostasis. The question that remains is how *mtHSP70* knockdown impacts mitochondria and ER contact. Although we did not explore this in our study, altered mitochondria and ER contact is seen in several tumor types as well as several neurodegenerative diseases, highlighting the importance of further research into this field.

## Methods

### Strains and Culture

AGD2797 (*unc-119*(*ed3*) III; uthSi71[*vha-6*p::mRuby::*xbp-1s*::*unc-54* 3’UTR, cb-*unc-119*(+)]) and AGD3156 (*unc-119*(*ed3*) III; uthSi65[*vha-6*p::ERss::mRuby::*ire-1a* (344-967aa)::*unc-54* 3’UTR cb-*unc-119*(+)] IV) were generous gifts from the Dillin lab (UC Berkeley). SJ4005 (zcls4[*hsp-4*p::GFP]), CL2070 (dvIs70[*hsp-16.2*p::gfp]), AM140(rmIs132 [*unc-54*p::Q35::YFP]), AM101 (rmIs110 [F25B3.3p::Q40::YFP]), GRU102 (gnaIs2 [*myo-2*p::YFP + *unc-119*p::Aβ1-42]), TU3401(uIs69 [pCFJ90 (*myo-2*p::mCherry) + *unc-119*p::*sid-1*]), OK814 (*nfyb-1* (*cu13*)), N2 wild-type worms and the rest of the strains used were obtained from Caenorhabditis Genetic Center (CGC). All worm strains were grown at 20C using standard methods as described previously^67^. Nematode Growth Media (NGM) agar plates containing OP50 *E. coli* stain were used for normal growth, while HT115 bacteria was used for RNAi feeding. All RNAi clones were obtained from the Ahringer library^68^.

### RNAi Assays and imaging analysis

Unless stated otherwise, HT115 containing RNAi sequences were cultured overnight in LB containing 100 ug/ml of carbenicillin. Cultures were then seeded onto NGM plates containing 1mM IPTG and stored in a blacked-out box to dry under dark conditions. The RNAi plates must be dried before use and long-term storage to ensure induction of T7 RNA polymerase activity. Worms were then transferred onto RNAi plates during their appropriate time points and grown at 20°C while remaining in the black box away from light. GFP reporter quantification was done by ImageJ. Worms were blindly picked under the LED dissecting microscope, then moved to the M805 Leica Fluorescent microscope to take the image. Briefly, 100 worms were treated per condition, and 6-10 worms were randomly picked under an LED dissecting microscope (blinded); then, worms were transferred to the fluorescent microscope room where they were put into sleep with 2mM levamisole solution. Then the fluorescent images were taken using M805. Then, individual worms were quantified with the ImageJ tools. Three technical repeats and three biological repeats are done for each condition.

### Tunicamycin Treatment

Worms were synchronized with the bleaching method and grown on OP50 *E. coli* until early adulthood (55 hours post-bleaching). Then, the worms were transferred to RNAi-containing or empty vector control plates. At 72 hours post-bleaching, worms were washed with M9 and treated with tunicamycin (25ug/ml) for four hours. After the tunicamycin treatment, worms were washed with M9 three to five times before transferring them back to fresh RNAi-containing or empty vector control plates. Animals were imaged on day 3 of the adult stage.

### Western Blots

Post RNAi treated worms were washed with M9 buffer followed by lysis in RIPA buffer (50 mM Tris-HCl pH 7.5, 150 mM NaCl, 1% NP-40, 0.1% SDS, 2 mM EDTA, and 0.5% sodium deoxycholate) with protease inhibitor cocktail (Roche) at their appropriate time points. The homogenized lysate was then centrifuged at 13,000RPM for 10 minutes at 4°C. The supernatant was then transferred into a new 1.5mL tube to determine protein concentration using BCA assay (Invitrogen). 20-40 ug of protein was loaded per lane in a pre-cast gel for SDS-PAGE (Bio-Rad). Gel was then transferred using the TurboBlot Transfer (Bio-Rad) followed by probing of phosphorylated eIF2α, a-tubulin, VCP and HSP-6. Quantification of bands was further carried out using ImageJ.

### Immunohistochemistry

Worms were fixed by following the previously described Buoin’s tube fixation ^69^ Briefly, 10 plates of worms with RNAi treatment were washed with M9 buffer and distilled water. Pelleted worms were fixed in Bouin’s MB fix solution, followed by liquid nitrogen to crack the cuticles. Permeabilized worms were then washed with BTB solution and shaking incubated in BTB solution. BTB solution was replaced with BT solution before incubating with fresh antibody buffer. Antibodies were incubated as described in the antibody buffer. Images were taken using a Nikon A1R confocal laser scanning system equipped with a CIF Plan-Apo 100X objective (NA 1.49, Nikon) and ORCA-Flash 4.0 CMOS camera (Hamamatsu). Primary antibodies: VCP (Abcam), and YFP (Roche). Secondary antibodies: sheep 488 and mouse 647 (Invitrogen)

### Paralysis Assay

Adult worms were synchronized using standard bleaching methods as described previously^70^. Worms were grown on standard NGM agar plates containing OP50 at 20C until day 1 adult. On day 1, 100 worms were transferred per RNAi condition onto NGM plates containing RNAi culture and transferred into the 25°C incubator. Worms were transferred onto fresh RNAi plates every other day, and paralysis of the worms was recorded every other day from day 1 to 5. Following day 5, the paralysis of the worms was recorded every day until all 100 worms were either dead or paralyzed. Worms that died due to external circumstances and of natural causes were censored during data recording to maintain consistency throughout the experiment. Data was recorded and analyzed using Excel and Prism.

### Body Bending Assay

Am101;Tu3401 worms were grown at 20°C on Nematode Growth Media (NGM) plates spotted with OP50 *E. coli*, then synchronized using the *C. elegans* standard bleaching method as mentioned previously^70^. On day 1 of adulthood, worms were transferred to NGM plates spotted with Empty Vector (EV), *hsp-6*, *hyl-1*, and *hyl-2* RNAi. They were incubated at 20°C (a typical incubation temperature for *C. elegans* growth) in the dark for the rest of the assay to prevent degradation of the IPTG that was inducing the RNAi. Worms were transferred to new RNAi plates every other day and progeny were removed on subsequent days as needed. Motility was measured on day 1 and day 5 of adult lifespan. To measure motility, worms were placed in a drop of M9 media and recorded for 30 seconds under a microscope. The number of body bends each worm performed within the 30 second timespan was manually counted and quantified with ImageJ.

### DAF28::GFP Assay

Daf-28::GFP transgenic worms were synchronized using standardized *C. elegans* bleaching method and eggs were plated on OP50 plates at 20°C to allow for normal development. On day 1 of adulthood, the worms were treated with RNAi targeting *hsp-6*, *xbp-1*, *ire-1* and EV (control). Worms were transferred to new RNAi plates on day 3, followed by imaging of the coelomocytes within the tail region on day 4. GFP intensity was recorded and quantified using ImageJ.

### Statistical analysis

Statistical analysis was performed as described in the figure legends. If not specified, paired student’s t-test is applied for comparing two groups, and one-way ANOVA with multiple comparisons is applied to for comparing multiple treatments to the control. All assays were repeated with at least three independent experiments. Data distribution was assumed to be normal but was not formally tested.

## Acknowledgment

We thank the Caenorhabditis Genetic Center and Shohei Mitani of the National BioResource Project for providing strains. We thank Dr. Daniel Garsin from University of Texas Health Science Center at Houston for sharing RNAi clones and other reagents. We thank Dr. Andrew Dillin and lab members for sharing the strains with us. We thank Jina Park, an English major Rice University student, for proofreading the manuscript. We thank Yelena Robinson, a Rice University student, and Tuckger Engels, a Medical student at the University of Texas Health Science Center at Houston (McGovern Medical School), for technical support.

This research was supported by the University of Texas Health Science Center at Houston (37516-12002), the Rising STARs program of UT systems (26532), and National Institutes of Health (NIH) K01 (NHLBI K01HL143111).

Fluorescence microscopy was performed at the Nikon Center of Excellence - Center for Advanced Microscopy, Department of Integrative Biology & Pharmacology at McGovern Medical School, UTHealth Houston.

The authors declare no competing financial interests.

## Author contributions

H-E Kim conceived the study. LA Tavizon performed ER secretion experiments and genetic crosses. R Hong performed paralysis and motility assays. J Park analyzed lifespan data with JJ Li. TI Moore analyzed the confocal images. RG Tharyan performed worm sorter experiments. JJ Li and N Xin carried out the rest of the experiments with the help of C Yang, and analyzed the data with input from H-E Kim. JJ Li and H-E Kim took the lead in writing the manuscript. All authors discussed the results and contributed to the final manuscript. H-E Kim was in charge of the overall direction. A Antebi supervised the project performed at the Max Planck Institute for Biology of Ageing, Cologne, Germany.

## Figure Legends

**Figure S1**. **a)** Inhibition of UPR^ER^ by *hsp-6* RNAi is regulated through the *dve-1* transcription factor. Animals were treated with tunicamycin as described above for 4 hours on day 1 of adulthood, followed by transfer onto RNAi plates, which targeted specific transcription factors within the mitochondria or cytosolic stress pathway. The animals were imaged at day 4 adult. **b)** Post-development mitochondrial stress through knockdown of different mitochondrial proteins has a different effect on the induction of UPR^ER^. Animals were treated with tunicamycin at L4, then were treated with the indicated RNAi or the empty vector control. Animals were imaged at day 4 adult. Graph shows mean+/-SD of four biological repeats, n>=8. **c)** Developmental mitochondrial stress by *cco-1* knockdown also does not affect UPR^ER^. UPR^ER^ (*hsp-4::gfp*) reporter strains in wild-type (N2) or *nfyb-1(cu13)* background were treated with control (luciferase) or *cco-1i* RNA from hatch, and GFP expression was measured on day 1 of adulthood with biosorter (left), and the intensity was normalized with the size of the worms (TOF: time of flight). Statistics determined by one-way ANOVA, ns: not significant, * p<0.5, ** p<0.01, Error bar shows mean± s.e.m.

**Figure S2. a)** *pek-1* mediates the suppression of UPR^ER^. Animals were treated with tunicamycin and the indicated RNAi as previously described (Figure 1). Graph shows mean+/-SD of three biological repeats, n>=6. **b)** Western blot of *daf-28*p::GFP and DAF-28::GFP shows that *hsp-6*, *ire-1* and *xbp-1* RNAi treatment does not decrease *daf-28* transcriptional and translational expression levels, suggesting that the reduction of *daf-28* exhibited in the coelomocytes in Figure 3f is most likely the result of a decrease in ER secretory function. **c)** Lifespan of the animals expressing polyQ35::YFP in their body wall muscle (AM140) with constitutively active UPR^ER^ in the whole-tissue by expressing *xbp-1s* (*myo-*3p::polyQ35::YFP; sur*-5*p::*xbp-1s*,*myo-2*p::tdTomato). Log-rank * p<0.05 for EV vs *hyl-2*.

**Figure S3. A working model of the proposed mechanism of MERSR.** Our findings suggest that the mitochondrial stress response (MCSR), triggered by HSP-6 knockdown, modulates UPR^ER^ signaling pathways through alterations in ceramide and cardiolipin levels. As misfolded proteins build up in the ER, the mitochondrial stress response curtails IRE-1 activation and turns-off the expression XBP1 target genes. Simultaneously, it promotes the PERK-dependent eIF2α phosphorylation, leading to a reduction in global protein translation. Consequently, as a newly discovered branch of MCSR, MERSR increases the ER stress threshold and the ER’s protein processing capacity, enhancing overall cellular proteostasis.

