## Supplemental Figures for "Unveiling the Intercompartmental Signaling Axis: Mitochondrial to ER Stress Response (MERSR) and its Impact on Proteostasis"

### Slide 1
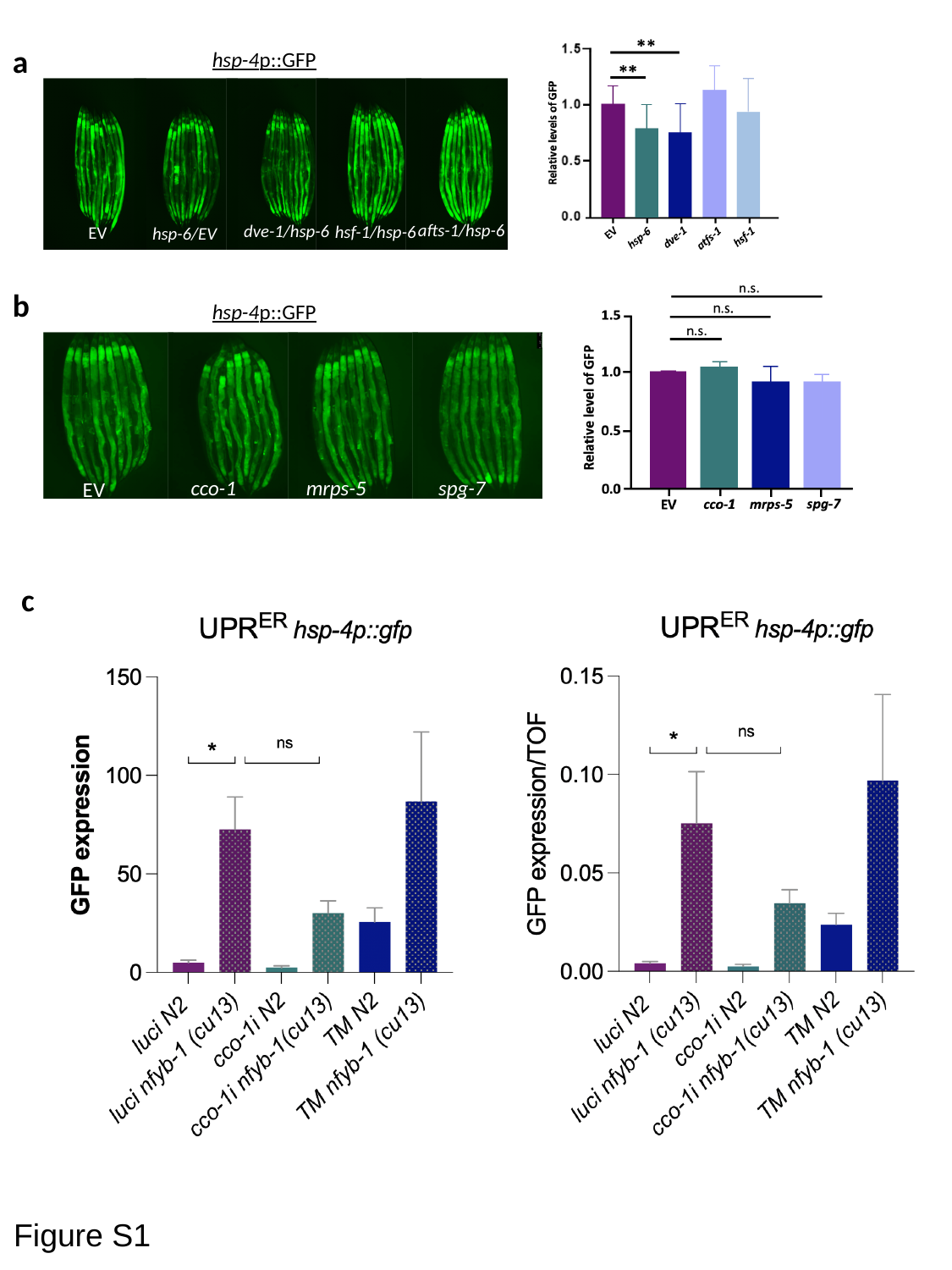

a
hsp-4p::GFP
afts-1/hsp-6
dve-1/hsp-6
hsf-1/hsp-6
EV
hsp-6/EV
b
hsp-4p::GFP
cco-1
mrps-5
EV
spg-7
c
Figure S1

### Slide 2
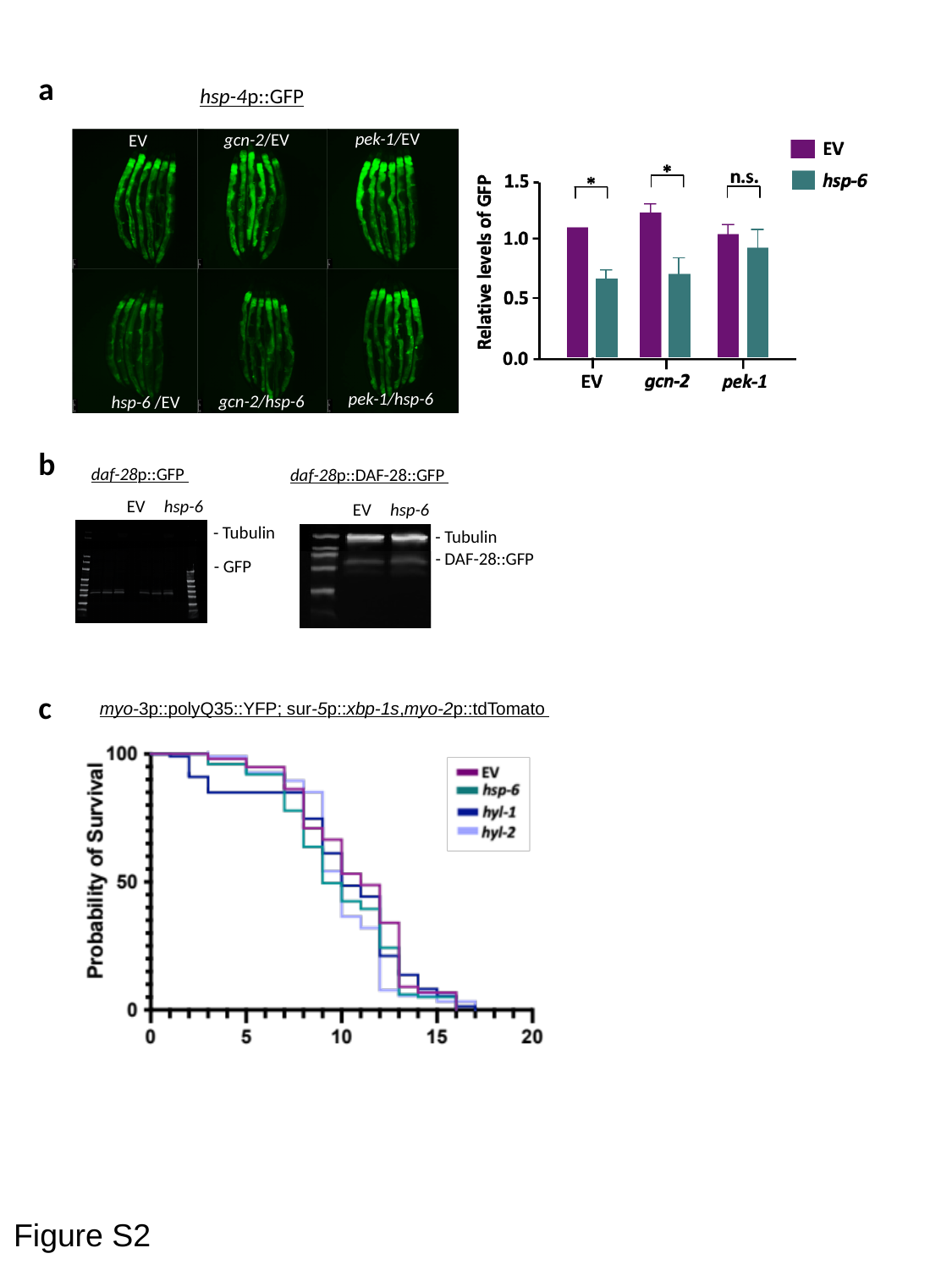

a
hsp-4p::GFP
pek-1/EV
gcn-2/EV
EV
pek-1/hsp-6
gcn-2/hsp-6
hsp-6 /EV
b
daf-28p::GFP
daf-28p::DAF-28::GFP
EV
hsp-6
EV
hsp-6
- Tubulin
- Tubulin
- DAF-28::GFP
- GFP
c
myo-3p::polyQ35::YFP; sur-5p::xbp-1s,myo-2p::tdTomato
Figure S2

### Slide 3
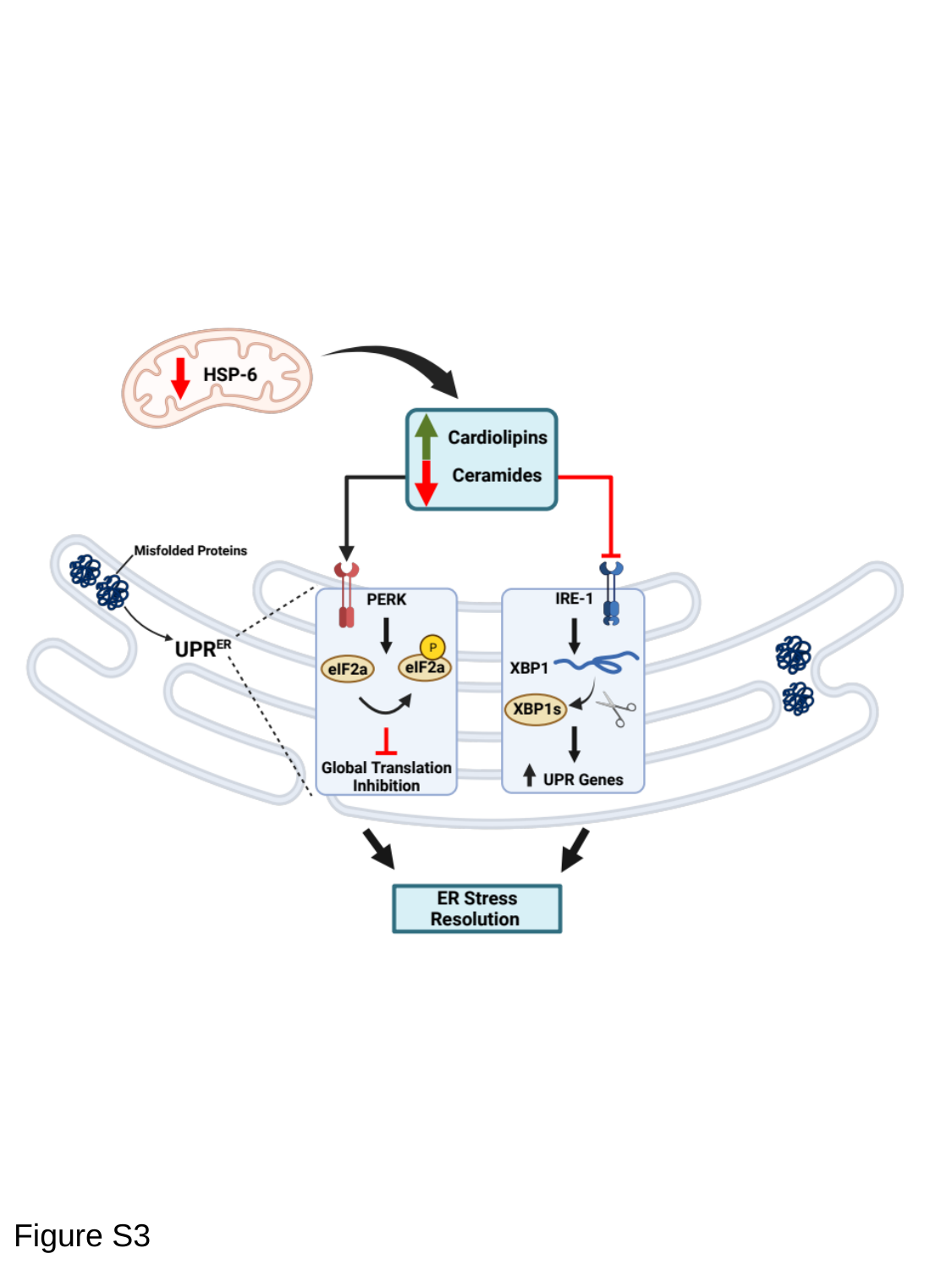

Figure S3
